# Multi-Scale, Patient-Specific Computational Flow Dynamics Models Predict Formation of Neointimal Hyperplasia in Saphenous Vein Grafts

**DOI:** 10.1101/624312

**Authors:** Francesca Donadoni, Cesar Pichardo-Almarza, Shervanthi Homer-Vanniasinkam, Alan Dardik, Vanessa Díaz-Zuccarini

## Abstract

Bypass occlusion due to neointimal hyperplasia (NIH) is among the major causes of peripheral graft failure. Its link to abnormal hemodynamics in the graft is complex, and isolated use of hemodynamic markers insufficient to fully capture its progression. Here, a computational model of NIH growth is presented, establishing a link between computational fluid dynamics (CFD) simulations of flow in the lumen, with a biochemical model representing NIH growth mechanisms inside the vessel wall. For all 3 patients analyzed, NIH at proximal and distal anastomoses was simulated by the model, with values of stenosis comparable to the computed tomography (CT) scans.

## 1. Introduction

Autogenous vein bypass is the most common technique for peripheral arterial revascularization for severe peripheral arterial diseases but is prone to develop neointimal hyperplasia (NIH), a leading cause of bypass failure.^1^ Both experimental studies and clinical observations suggest that one of the factors destabilizing the remodeling process is a lower level of wall shear stress on the arterial wall,^2^ and numerous computational fluid dynamics (CFD) studies of blood flow have used shear stress indices - including Time-Averaged Wall Shear Stress (TAWSS), and Oscillatory Shear Index (OSI), for example - as markers to identify potentially problematic areas of vascular remodeling (**Supplementary Table I**).^3,4^

We hypothesized that both low and oscillatory levels of shear stress should be considered simultaneously when assessing the proclivity of a certain region in bypass grafts to develop NIH,^5,6^ and we simulated NIH progression using a multi-scale computational framework that we previously developed,^7^ comparing our results to a patient-specific clinical dataset (obtained with the patients’ informed consent for research and publication).

## 2. Methods

The computational framework was informed by patient-specific imaging data, including anatomical images and hemodynamic markers. Doppler scans immediately after surgery and CT scans at 8, 19 and 24 months after surgery (for patients 1-3, respectively) were obtained from routine clinical studies after approval from the institutional review board (IRB) (approval number AD0009, Veteran Affairs Connecticut Healthcare System, West Haven, CT, USA). CT scans immediately after surgery were not available, as the data was acquired retrospectively from selected patients that received standard of care with regular postoperative surveillance. Postoperative surveillance, at the institution where data was collected, is only performed via duplex ultrasound, as it is noninvasive, does not require nephrotoxic dye, is reproducible, and correlates with outcomes as documented with extensive literature, and as such, CT scans are only obtained when the duplex ultrasound suggested an abnormality and additional anatomic information is required. In order to overcome these limitations, a ‘baseline’ configuration (representing the vein-graft conditions right after implantation) was obtained by processing the images and ‘virtually removing’ regions of NIH growth, well in line with other work in the literature.^8–15^

Data required deidentification despite IRB approval in compliance to VA requirements for patients’ privacy.

CFD analyses were performed as described in a previous publication.^7^ A non-Newtonian Carreau-Yasuda model was used for blood viscosity, with parameters reported in previous studies.^16^ For comparison, simulations were also run with a Newtonian model (viscosity of 0.035 dynes∙s /cm^2^). Inflow conditions were described by flow curves obtained from Doppler scans. These were applied first with a flat profile, and with a parabolic profile in a different set of simulations for comparison. Boundary conditions at the outlets were implemented via two-element Windkessel models of the external vasculature (**Supplementary Figure 1**), tuned to patient-specific data on a 0D model (**Supplementary Table II)**.

Simulation results were processed using ANSYS CFD Post (Ansys, Inc.). Hemodynamic stress indices linked to vascular remodeling, specifically TAWSS and a term encompassing low shear together with oscillations - the highly oscillatory, low magnitude shear (HOLMES) - (**Supplementary Table I**) were extracted at each node and imported into a mathematical model of NIH progression, described in **Supplementary Figure 2**. The output of the simulation model for each patient is the predicted (calculated) values of NIH growth along the graft, following the same blueprint described in our previous work,^7^ and summarized in **Supplementary Table III**. In a nutshell, based on the ‘base’ configuration for each patient (a vein-graft free of NIH, representing conditions just after implantation), CFD analyses are coupled to a mathematical (time-dependent) model of NIH growth that takes into account smooth muscle cells and collagen turnover, growth factors and NO production. This is a dynamic simulation process that captures the transformation of the vein-graft due to NIH and mimics the evolution of the disease for each patient, in time. A diagram of the different cases is presented in **Supplementary Figure 3**.

## 3. Results

Hemodynamic analyses were performed on all three bypass geometries (Figure 1). As shown in Figure 2, at the proximal and distal anastomosis where NIH developed, low values of TAWSS (<0.5 Pa) did not always correspond to regions of NIH progression. Similarly, high values of OSI did not always fully capture the areas of NIH progression either. HOLMES was the only index able to capture all regions of NIH progression (Figure 2a).

**Figure 1:**
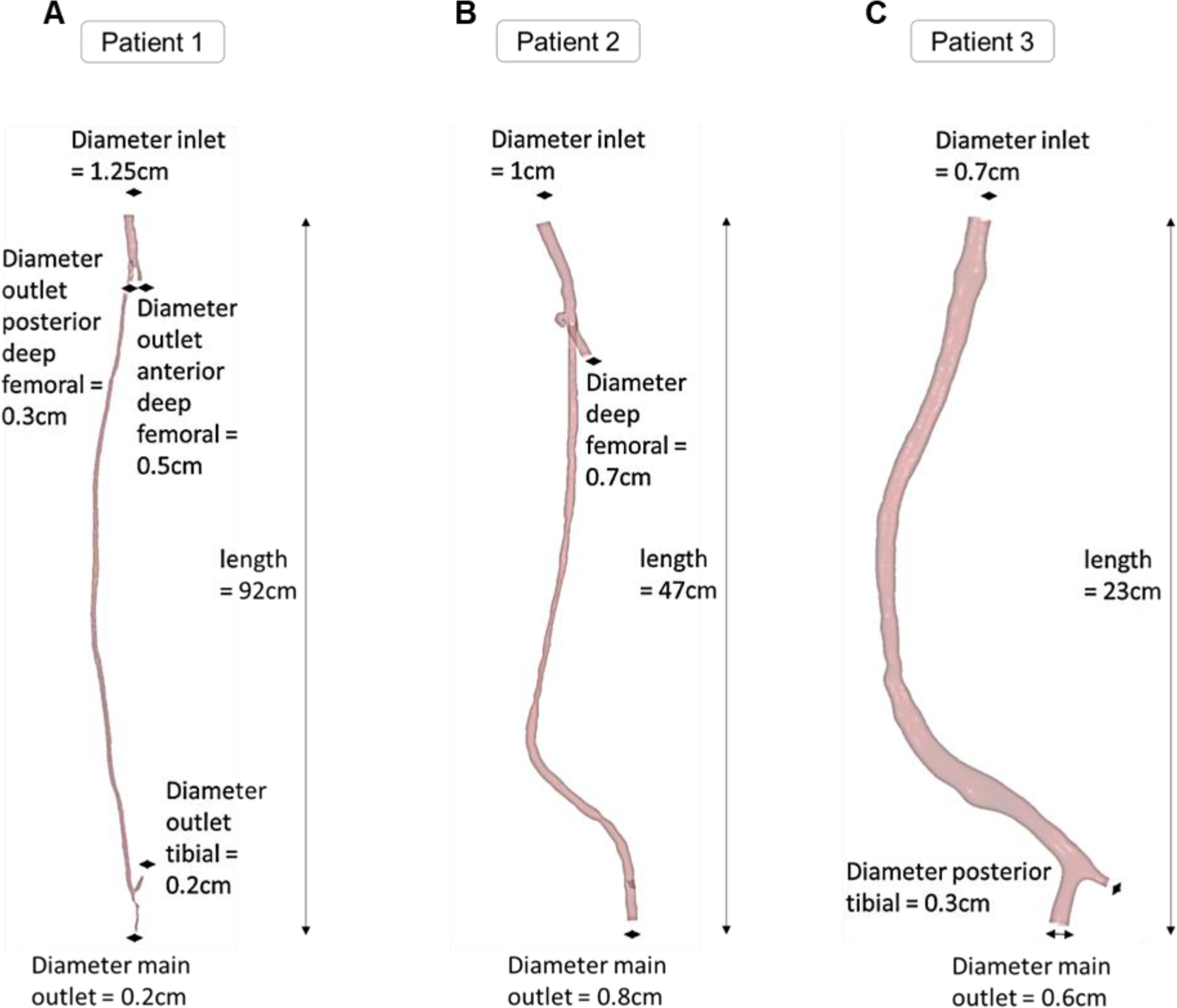
Summary of the geometrical characteristics of patients analyzed in the study. A. Patient 1: Femoro-distal bypass; B. Patient 2: Femoro-popliteal bypass; C. Patient 3: Femoro-popliteal bypass.

**Figure 2:**
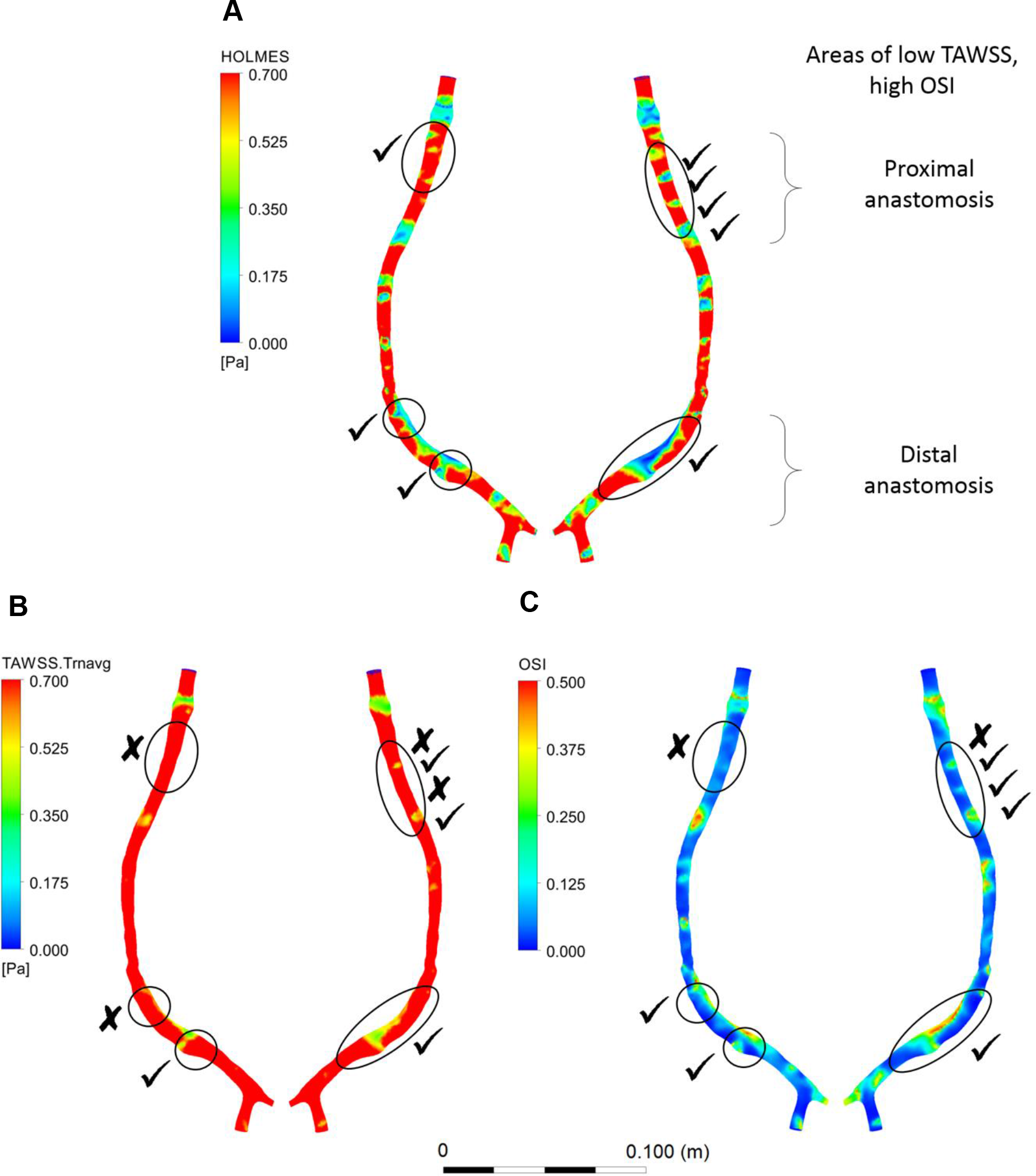
Example of contour plots of hemodynamic indices obtained for patient 3. A. Simulation for HOLMES; B. Simulation for TAWSS; C. Simulation for OSI. The ‘tick’ (✔) boxes refer to areas of NIH actually captured by CFD simulations. It can be seen that in the case of TAWSS and OSI many remodeled regions are not indicated (identified by x).

Percentage differences between the initial and remodeled lumen areas were calculated and are reported in **Supplementary Table IV** and displayed in Figure 3. In all the cases simulated, predicted stenoses were confirmed by the CT scan data, and only one of the bypasses showed a considerable amount of NIH in the mid-region of the graft that was not captured by the model since this area coincided with an area of pre-existing vein graft stenosis, another cause of NIH.^17^

**Figure 3:**
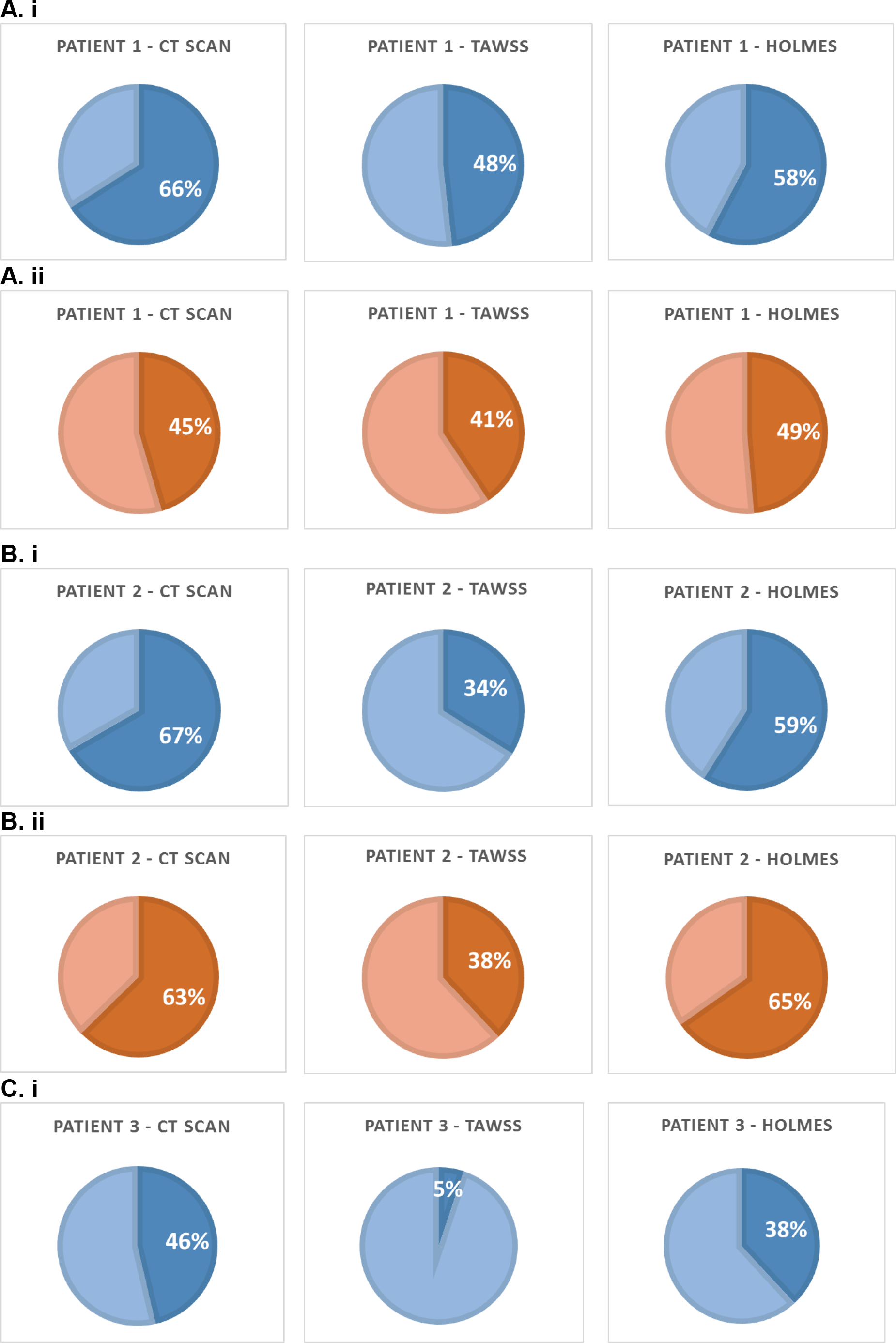
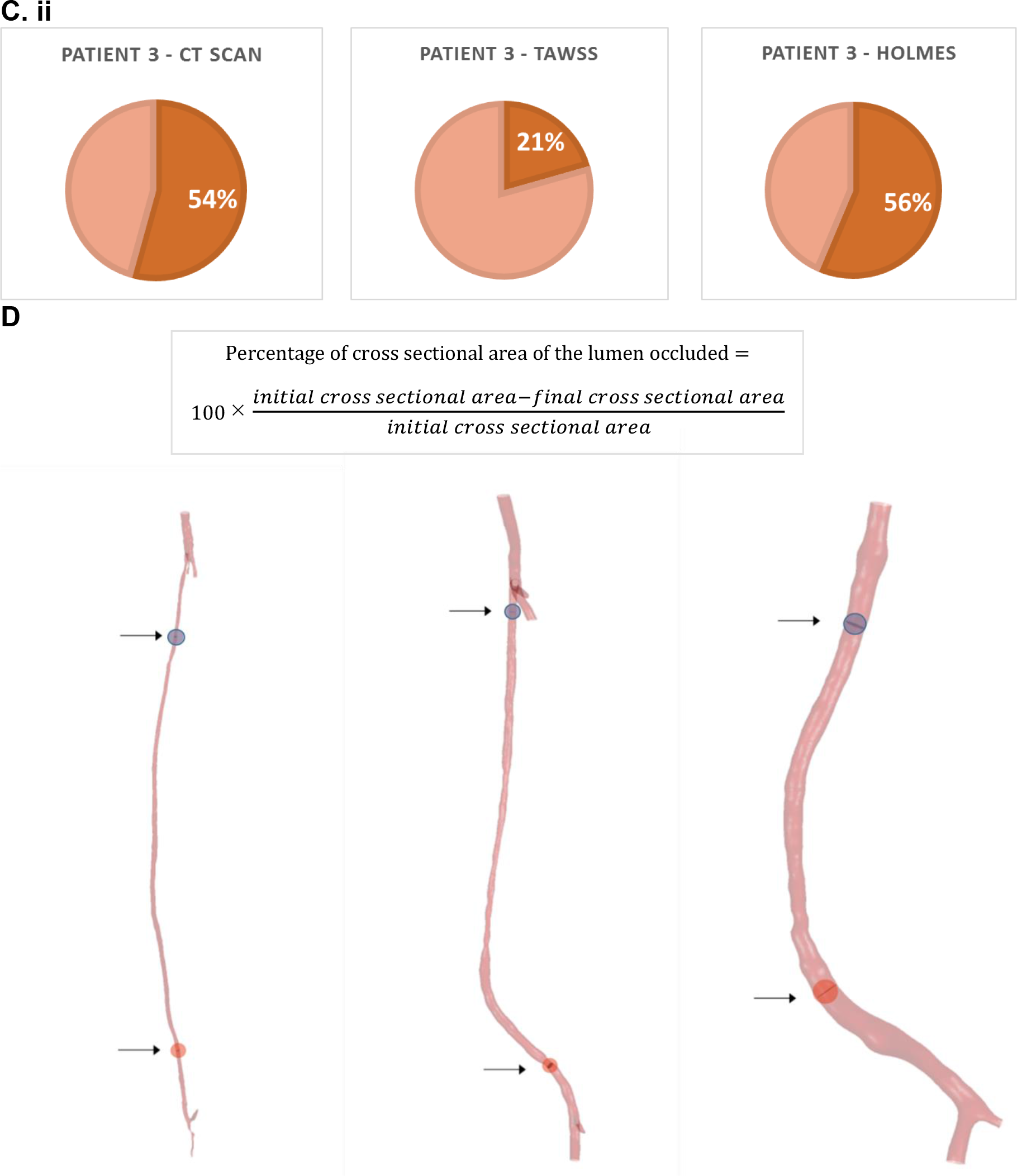
results of the simulations in % cross sectional area occupied by NIH at the most severely affected locations after the time of NIH development for each patient, using the non-Newtonian viscosity model and the two different wall shear stress indices (TAWSS and HOLMES). A.i. Patient 1 - proximal anastomosis; A.ii. Patient 1 - distal anastomosis; B.i. Patient 2 - proximal anastomosis; B.ii. Patient 2 - distal anastomosis; C.i. Patient 3 - proximal anastomosis; C.ii. Patient 3 - distal anastomosis; D. Calculations of occlusion at the locations most severely affected by NIH.

**Supplementary Figure 4** and **5** summarize simulation results of the mathematical model used to describe cell proliferation and collagen turnover.^7^ To the authors’ best knowledge, values for collagen and smooth muscle cell composition *in vivo* are not reported in literature; however, the figures show an increasing number of smooth muscle cells and collagen in areas of low wall shear and high oscillations, which agrees with findings widely reported.^18^

Finally, analysis of graft geometry was performed to measure curvature, torsion, and tortuosity using the Vascular Modeling Toolkit (VMTK)^19,20^ (www.vmtk.org). Values of torsion and tortuosity were calculated at specific locations of interest and presented the highest differences among NIH prone locations (up to 99.3%, **Supplementary Table V**). The highest average torsion was found in the distal segment of patient 1 (0.12 mm^−1^), where the average value of TAWSS was 0.69 Pa and HOLMES was 0.34 Pa.

## 4. Discussion

These results highlight the impact of two different measures of wall shear stress, TAWSS and HOLMES, and the importance of the interaction between TAWSS and OSI in NIH progression (**Supplementary Figure 6**). In all cases, the simulation model *correctly predicted* areas of NIH growth, with values that were similar to the stenoses observed in the CT scans when using the HOLMES index, with a maximum discrepancy (presented as % area) of 8% between stenosis values observed in patients 1-3 when compared to CT scans (Figure 3). When using TAWSS, not all NIH-stenotic regions are predicted and for those that are, the amount of luminal narrowing is consistently underestimated and sometimes by a significant amount - as in the case of patient 3 - with a reported difference in terms of NIH growth area of 41% (Figure 3). This suggests that TAWSS is an unreliable metric to estimate both plaque location and the degree of stenosis in vein-grafts.

CFD has been used to analyze the hemodynamics of grafts for multiple cardiovascular procedures, such as endovascular repair, ^21^ coronary stenting ^22^ and AVF. ^23^ Much less research has been reported on mechanisms of failure of peripheral bypasses, with most reported work focused on design optimization.^24^ This preliminary study of 3 patients couples CFD analyses with a model of smooth muscle cells and collagen turnover, including growth factor and NO production.^7^ The results show a change from an initial, predominantly homogeneous distribution of smooth muscle cells and collagen at day 146 to a localized area of growth corresponding to areas in which the graft experienced low shear and oscillations (**Supplementary Figure 4 and 5**). In addition, our analysis shows that morphometric indicators alone might not be enough to identify successful grafts. For instance, torsion has been observed to be one of the key factors affecting large vessel hemodynamics,^25^ and our results agree with these findings. However, the results presents too high variability and thus are difficult to generalize.

### 4.1. Limitations

Multiple factors affect NIH remodeling, such as pre-existing valve lysis, as demonstrated by the results for patient 2, which should be included in future simulation frameworks. The framework could also be improved by considering additional biological mechanisms related to growth factors, extracellular matrix components, and endothelial cells, all of which have been shown to influence NIH.^1^ Patient-specific biomarkers such as C-reactive protein, inflammatory cytokines, and adhesion molecules^26,27^ also correlate with severity of NIH. External factors such as smoking and medication use, as well as other clinical conditions such as hypertension could play a role in the outcome and should be considered in future developments of the model. The voxel size in the CT scans is close to the order of magnitude of the geometries considered and this might have an impact in the results. Finally, the model might also be limited by the assumption of rigid arterial walls and the impact of this assumption needs to be verified through further analyses.

## 5. Conclusions

In the application of a multi-scale model of NIH on three different patient-specific cases, the HOLMES index was the best predictor for the location of NIH progression that corresponded to developing stenoses identifiable in CT scans. Results also suggest that TAWSS is an unreliable predictor of NIH growth. The analysis demonstrated how multiscale modeling may play an important role in the post-revascularization management of PAD patients, and specifically, in delineating those at risk of developing NIH.

## 6. Acknowledgements

The authors gratefully acknowledge support by the Leverhulme Trust Senior Research Fellowship “Exploring the Unknowable Using Simulation: Structural Uncertainty in Multiscale Models” (Fellowship number RF-2015-482). This work was supported by the resources and the use of facilities at the Veterans Affairs Connecticut Healthcare System (West Haven, CT). The authors gratefully acknowledge funding from the Wellcome/EPSRC Centre for Interventional and Surgical Sciences, made possible by Wellcome/EPRSRC grant 203145Z/16/Z

