## Supplementary data for "Multi-Scale, Patient-Specific Computational Flow Dynamics Models Predict Formation of Neointimal Hyperplasia in Saphenous Vein Grafts"

### 1. Supplementary Tables

*Supplementary Table I: A summary of shear stress indices used in this study*

| Index | Formula | Meaning and Relevance in cardiovascular disease |
|---|---|---|
| **TAWSS** | $\frac{1}{T}\int_0^T \lvert\vec{\tau}_w\rvert dt$ <br> where T = time and $\tau_\omega$ = wall shear stress. | A measure of wall shear stress for one cardiac cycle: values lower than 0.5 Pa co-localize with areas of neointimal thickening. |
| **OSI** | $0.5\left(1-\frac{\left\lvert\int_0^T \vec{\tau}_w\, dt\right\rvert}{\int_0^T \lvert\vec{\tau}_w\rvert dt}\right)$ | Used to identify areas where the flow deviates most from its average direction, that is where it oscillates most. |
| **HOLMES** | $TAWSS\,(0.5 - OSI)$ | A modified TAWSS, highlighting regions of low shear and modulated by high oscillations. |

*Supplementary Table II: summary of the parameters used for the 2-element Windkessel models. R units: (mmHg s ml$^{-1}$); C units: (ml mmHg$^{-1}$)*

| | $R_1$ | $R_2$ | $R_3$ | $R_4$ | $C_1$ | $C_2$ | $C_3$ | $C_4$ |
|---|---|---|---|---|---|---|---|---|
| Patient 1 | 12.5 | 9.2 x10$^{2}$ | 1.5 x10$^{3}$ | 1.3 x10$^{3}$ | 2x10$^{-2}$ | 1.5 x10$^{-2}$ | 1.5x10$^{-2}$ | 1.5x10$^{-2}$ |
| Patient 2 | 25 | 2 x10$^{2}$ | | | 2.78 x10$^{-2}$ | 3.02 x10$^{-2}$ | | |
| Patient 3 | 13.9 | 1.89 x10$^{2}$ | | | 2 x10$^{-2}$ | 1.72 x10$^{-2}$ | | |

*Supplementary Table III: overview of the model characteristics*

| | | | Patient 1 | Patient 2 | Patient 3 |
|---|---|---|---|---|---|
| | | | **Patient 1** | **Patient 2** | **Patient 3** |
| **CFD model** | Mesh elements | | 1.6M | 380k | 200k |
| | Mesh sensitivity analysis | Coarse mesh | 700k | 120k | 70k |
| | | Medium mesh | 1.6M | 380k | 200k |
| | | Refined mesh | 2.6M | 600k | 500k |
| | | % difference in velocity at outlets between refined and medium mesh (<2% required for mesh convergence) | 1.6% | 0.9% | 1.9% |
| | Assumptions | • Non-Newtonian model of viscosity (Carreau-Yasuda)<br>• Flat profile at the inlet<br>• 2-element Windkessel models at the boundaries | | | |
| | Governing equations | Navier-Stokes (incompressible)<br>$\frac{\partial \boldsymbol{u}}{\partial t} + \nabla \boldsymbol{u} \cdot u - \nu \Delta \boldsymbol{u} + \nabla p = \boldsymbol{f}$<br>$\nabla \cdot \boldsymbol{u} = 0$<br>Where<br>u = velocity<br>p = pressure<br>$\nu$ = kinematic viscosity | | | |
| **NIH progression model** | Input | TAWSS or HOLMES | | | |
| | Output | Neointimal hyperplasia volume (*)<br>$V_i = (S_i + Q_i) \times \rho_s^{-1} + C_i \times \rho_c^{-1}$<br>Where<br>i = suffix referring to the intimal layer<br>S = smooth muscle cells<br>Q = quiescent cells<br>C = collagen<br>$\rho_s$ = cell density [cells/$m^3$]<br>$\rho_c$ = collagen density [cells/$m^3$]<br><br>*Neointimal hyperplasia volume is calculated based on a system of equations described in [Donadoni F, Pichardo-Almarza C, Bartlett M, Dardik A, Homer-Vanniasinkam S, Díaz-Zuccarini V. Patient-Specific, Multi-Scale Modeling of Neointimal Hyperplasia in Vein Grafts. Front Physiol. 2017 Apr 18;8:226]. | | | |

*Supplementary Table IV: Results from the CT scans and simulations for the three patients (% occlusion)*

| | | CT scans | Plug inflow - TAWSS | Parabolic inflow - TAWSS | Non-Newtonian viscosity - TAWSS | Plug inflow - HOLMES | Parabolic inflow - HOLMES | Non-Newtonian viscosity - HOLMES |
|---|---|---|---|---|---|---|---|---|
| Patient 1 | Proximal | 66.0 | 49.8 | 49.8 | 48.2 | 58.4 | 58.4 | 57.7 |
| | Distal | 48.9 | 43.5 | 43.5 | 42.8 | 50.3 | 50.3 | 50.0 |
| Patient 2 | Proximal | 66.7 | 44.2 | 44.3 | 33.9 | 66.1 | 66.2 | 58.9 |
| | Distal | 62.7 | 47.8 | 47.8 | 38.0 | 69.0 | 69.0 | 65.3 |
| Patient 3 | Proximal | 46.3 | 11.3 | 7.3 | 5.1 | 39.2 | 39.9 | 38.2 |
| | Distal | 54.3 | 25.2 | 27.0 | 20.6 | 56.1 | 54.7 | 56.3 |

*Supplementary Table V: Morphometric analysis of the three grafts*

| | Curvature (mm-1) | Torsion (mm-1) | Tortuosity |
|---|---|---|---|
| **Patient 1 proximal** | 0.034589 | 0.001381 | 0.011302 |
| **Patient 1 distal** | 0.026341 | 0.11598 | 0.012554 |
| **Patient 2 proximal** | 0.027653 | 0.101641 | 0.011055 |
| **Patient 2 distal** | 0.029005 | 0.0444 | 0.121906 |
| **Patient 3 proximal** | 0.034212 | 0.008087 | 0.012636 |
| **Patient 3 distal** | 0.037476 | 0.000842 | 0.051892 |

### 3. Supplementary Figures

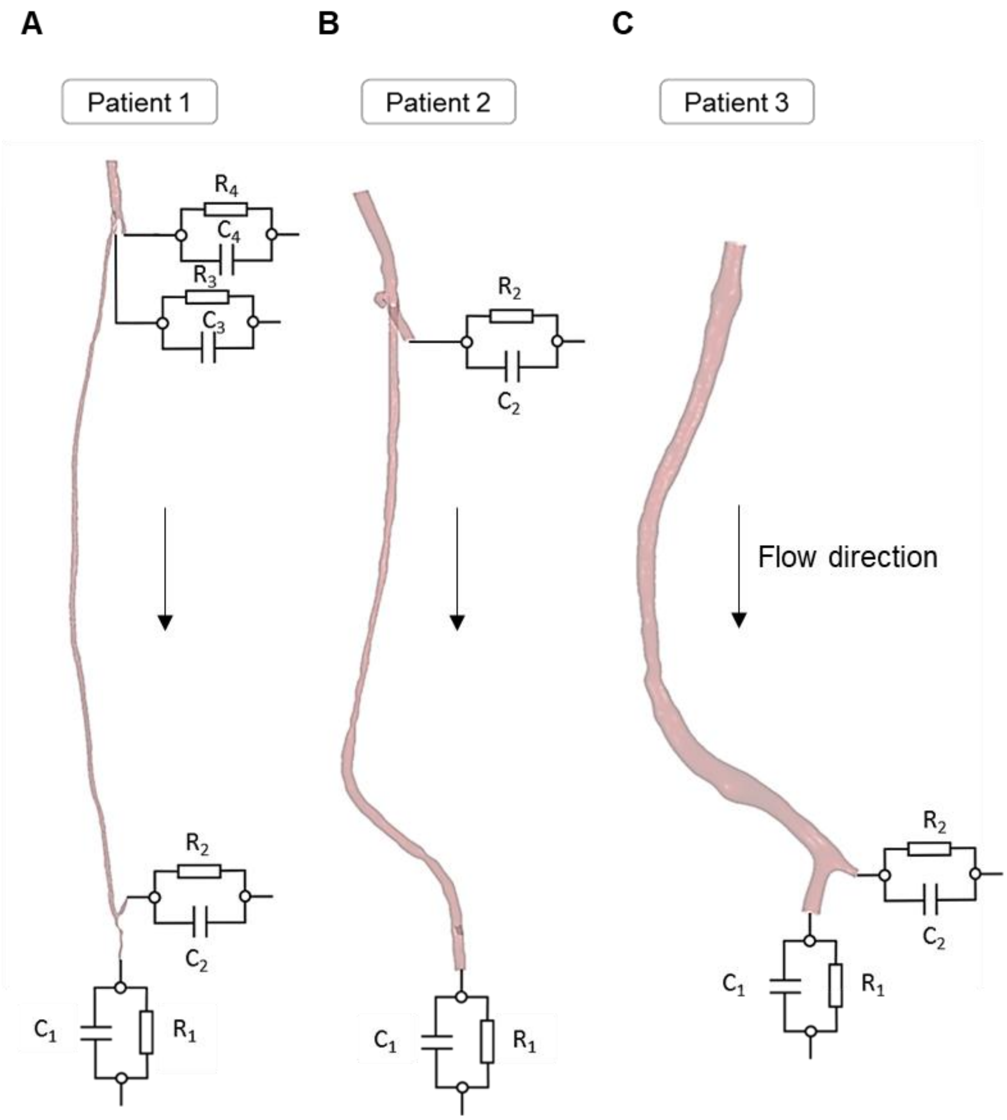


*Supplementary Figure 1: Boundary conditions at the outlets were modeled using RC elements. A. Patient 1: Femoro-distal bypass; B. Patient 2: Femoro-popliteal bypass; C. Patient 3: Femoro-popliteal bypass.*

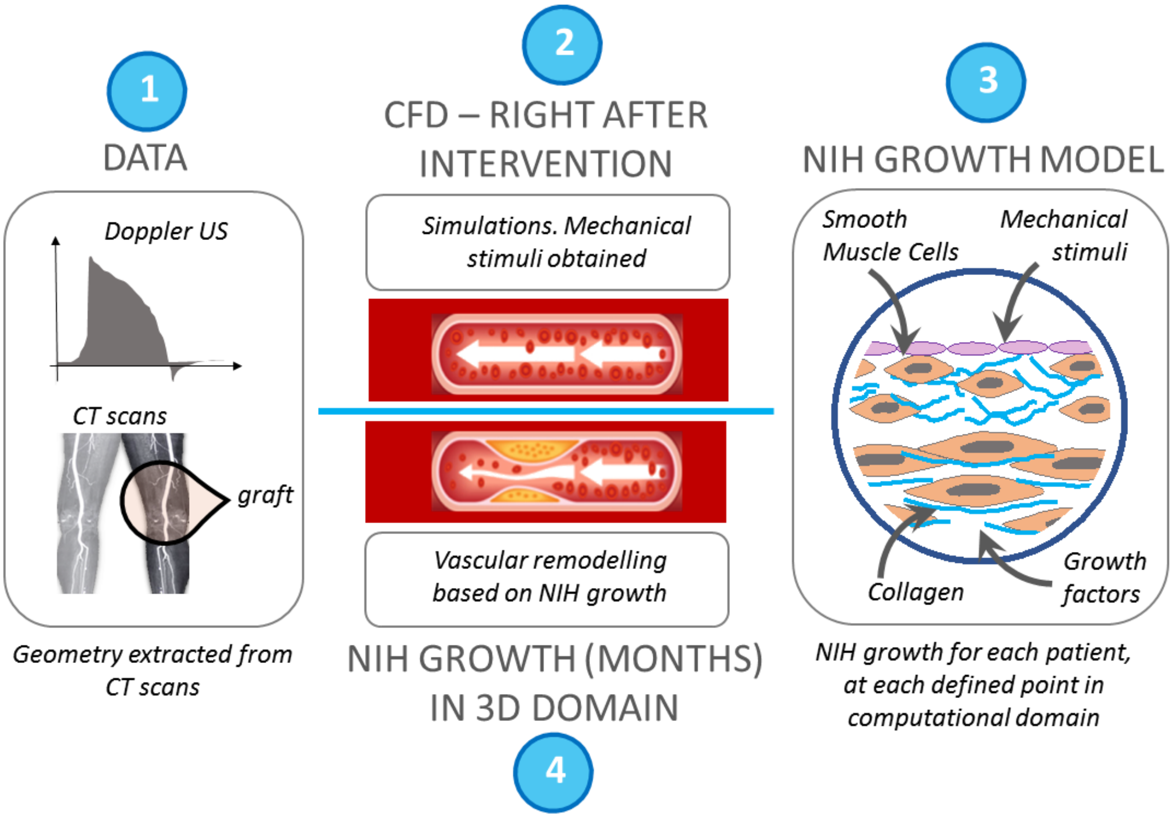


*Supplementary Figure 2: A schematic diagram of the framework for simulating NIH progression. 1. Clinical data is extracted for each patient (e.g. CT scans, Doppler ultrasound) and processed for CFD simulations; 2. CFD simulations are performed right after the intervention obtaining mechanical stimuli and calculating hemodynamic stress indexes (e.g. TAWSS, OSI, HOLMES); 3. Simulations of a biochemical model of NIH growth and progression; 4. Simulations of NIH growth for a long time span in the 3D domain.*

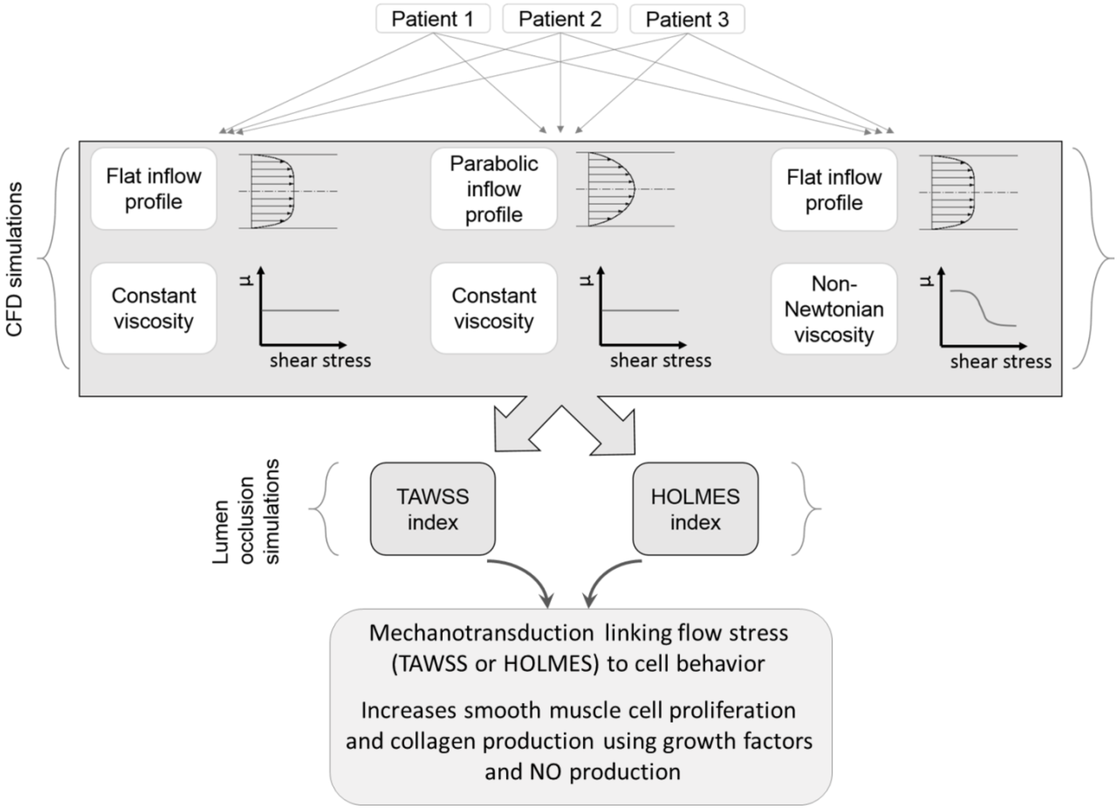


*Supplementary Figure 3: Overview of interactions: A complex interaction between mechanical stimuli on the wall affect cell behavior, with smooth cells and collagen proliferating due to growth factors and NO production. The result is the development of NIH and vascular remodeling. Details of the model are published in [Donadoni F, Pichardo-Almarza C, Bartlett M, Dardik A, Homer-Vanniasinkam S, Díaz-Zuccarini V. Patient-Specific, Multi-Scale Modeling of Neointimal Hyperplasia in Vein Grafts. Front Physiol. 2017 Apr 18;8:226].*

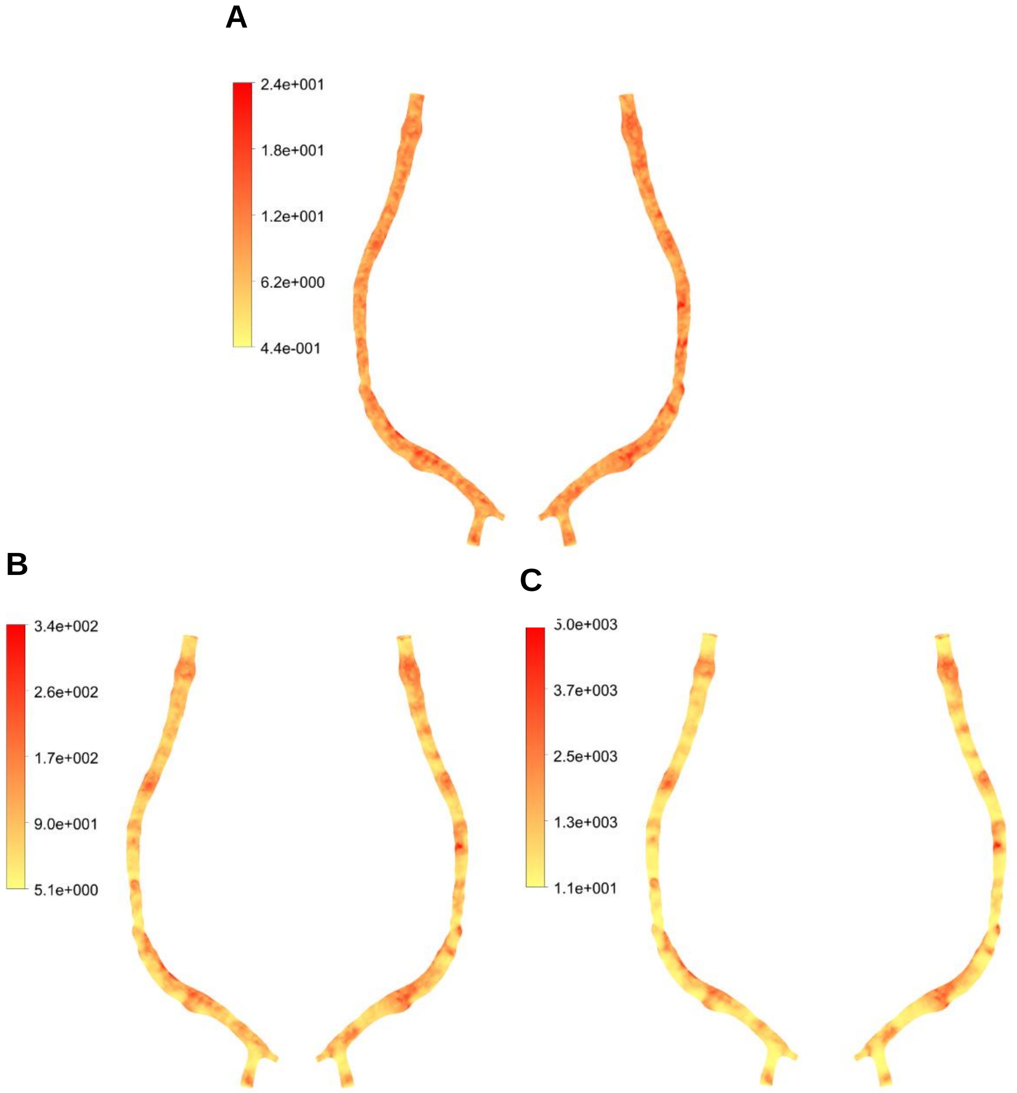


*Supplementary Figure 4: Smooth muscle cells per baseline value (363 cells/mm$^2$) in the intimal layer of the bypass at different time points during the NIH phase (Patient 3, non-Newtonian case using the HOLMES index). A. Snapshots at: 146 days after intervention; B. Snapshot at 438 days after intervention; C. Snapshot at 730 days after the intervention.*

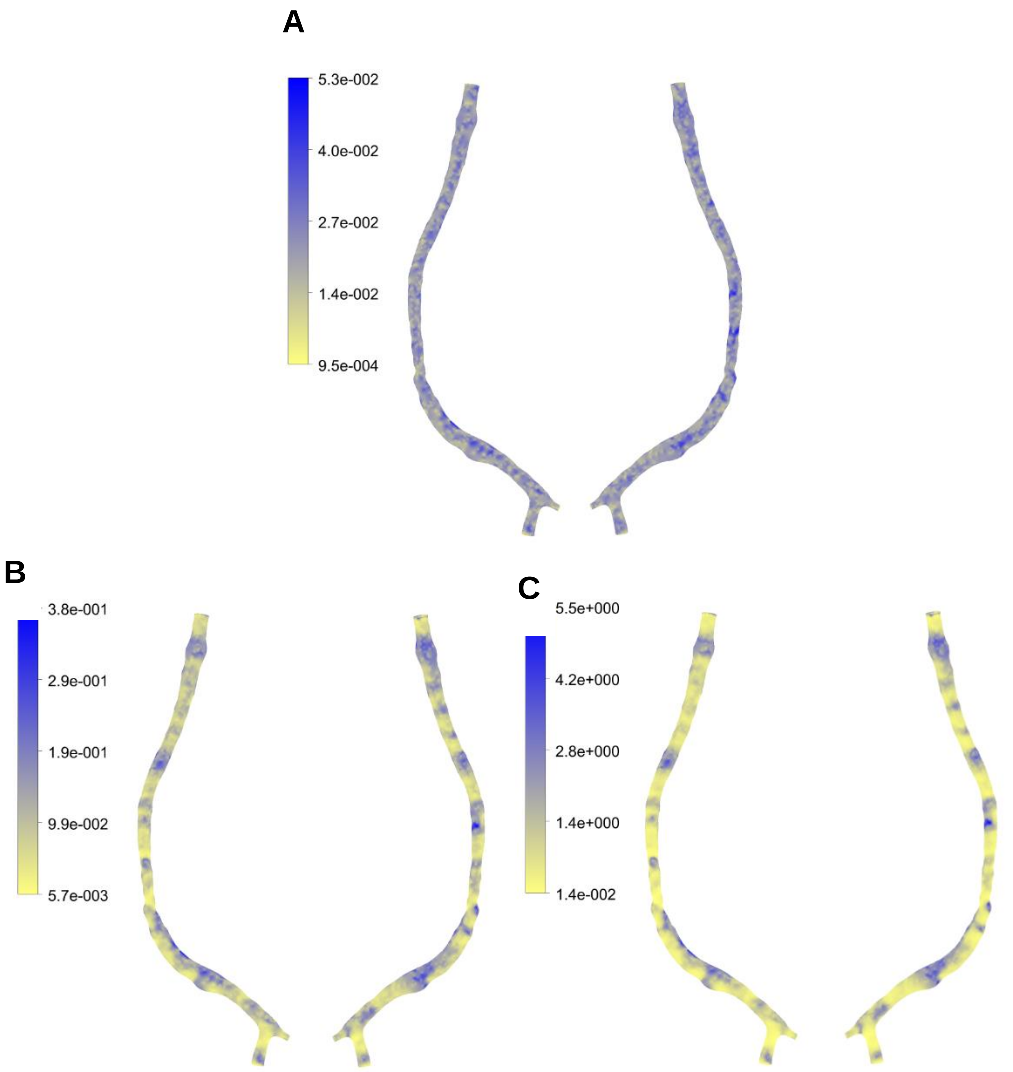


*Supplementary Figure 5: Evolution of collagen per baseline value (1.67 $\mu g/mm^2$) in the intimal layer of the bypass at different time points during the NIH phase (Patient 3, non-Newtonian case using the HOLMES index). A. Snapshots at: 146 days after intervention; B. Snapshot at 438 days after intervention; C. Snapshot at 730 days after the intervention. Snapshots presented at: 146, 438 and 730 days after the intervention.*

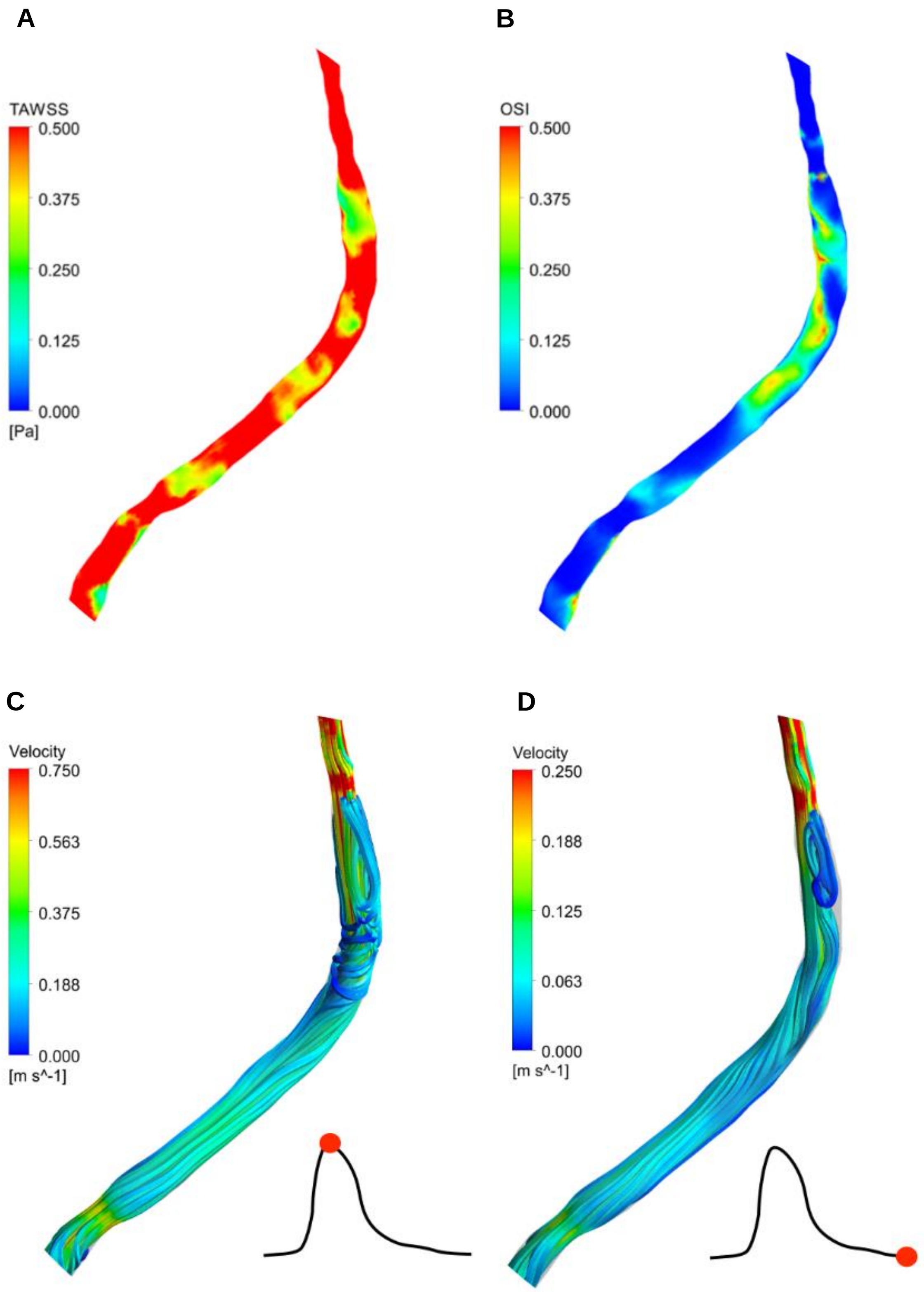


*Supplementary Figure 6: Contour plots of TAWSS and OSI and velocity streamlines at peak systole and end diastole in patient 2. A. TAWSS; B. OSI; C. Velocity streamlines at peak systole; D. Velocity streamlines at end diastole.*
